# Large-scale examination of neuropsychiatric, cognitive and cardiovascular phenotypic associations with 15q11.2 BP1-BP2 deletion in ∼500,000 UK Biobank individuals

**DOI:** 10.1101/722504

**Authors:** Simon G. Williams, Apostol Nakev, Hui Guo, Simon Frain, Gennadiy Tenin, Anna Liakhovitskaia, Priyanka Saha, James R. Priest, Kathryn E. Hentges, Bernard D. Keavney

**Affiliations:** Division of Cardiovascular Sciences, School of Medical Sciences, Faculty of Biology, Medicine and Health, Manchester Academic Health Science Centre, University of Manchester, Manchester, UK; Division of Population Health, Health Services Research and Primary Care, School of Health Sciences, Faculty of Biology, Medicine and Health, University of Manchester, Manchester, UK; Department of Pediatrics, Stanford University, Stanford, CA, USA; Division of Evolution and Genomic Sciences, School of Biological Sciences, Faculty of Biology, Medicine and Health, University of Manchester, Manchester, UK

**Author notes:** Corresponding author (BDK).

## Abstract

**Background:** Deletion of a non-imprinted 500Kb genomic region at chromosome 15q11.2, between breakpoints 1 and 2 of the Prader-Willi/Angelman locus (BP1-BP2 deletion) has been associated in previous studies with phenotypes including developmental delay, autism, schizophrenia and congenital cardiovascular malformations (CVM). The deletion has a low baseline population prevalence and large-scale data regarding the magnitude of these associations and any milder effects on cognition phenotypes, in populations not selected for disease, are limited.

**Methods:** Using the UK Biobank (UKB) cohort of ∼500,000 individuals, we identified individuals with neuropsychiatric and CVM diagnoses and investigated their association with deletions at the BP1-BP2 locus. In addition we assessed the association of BP1-BP2 deletions with cognitive function and academic achievement in individuals with no previous diagnosis.

**Results:** Cases of neurodevelopmental and CVM disease had an increased prevalence of the deletion compared to controls (0.68%; OR=1.84 [95% CI 1.23 – 2.75]; p=0.004 and 0.64%; OR=1.73 [95% CI 1.08 – 2.75]; p=0.03 respectively). Excluding participants diagnosed with neurodevelopmental or neuropsychiatric disease, deletion carriers had worse scores in four tests of cognitive function, and while 32.8% of UKB participants without BP1-BP2 deletion had a university or college degree as their highest educational qualification, only 22.8% of deletion carriers achieved this (OR 0.57 [95% CI 0.51-0.64]; p=5.60E-22).

**Conclusions:** We conclude that BP1-BP2 deletion has an appreciable population prevalence with important life-course impacts on undiagnosed carriers. These data are of potential utility in deciding the circumstances under which clinical testing for BP1-BP2 deletion may be helpful.

**Key messages:**

- Deletions at chromosome 15q11.2 between breakpoints 1 and 2 (BP1-BP2), which encompass four genes, have been associated with developmental delay, autism, schizophrenia and congenital cardiovascular malformations.
- Here, we use the largest cohort studied to date: the UK Biobank cohort of ∼500,000 individuals, to explore the association of this deletion with neuropsychiatric and cardiovascular phenotypes as well as its effects on cognitive function and academic achievement.
- We find an appreciable population prevalence of the deletion with these phenotypes and demonstrate reduced cognitive function and lower academic achievements in those individuals with no prior diagnosis.

## Introduction

Copy number variants (CNVs) are a form of genomic structural variation consisting of deletions or duplications of genetic material. These variations may encompass the coding regions of multiple genes. CNVs are known to be common, possibly accounting for as much as 13% of the human genome ^1^. Studies have concluded that CNVs which occur commonly in the population are unlikely to contribute significantly to human disease ^2^, while in some cases the presence of CNVs may confer a functional advantage ^3^. However, many genes are sensitive to copy number and alterations can lead to dosage imbalance ^4^. Consequently many rare CNVs have been identified that are associated with a variety of diseases including schizophrenia ^5, 6^, autism ^7^, developmental disorders ^8, 9^ and congenital cardiovascular malformations (CVM) ^10^.

15q11.2 BP1-BP2 deletions occur between breakpoints 1 and 2 (BP1-BP2) on the long arm of chromosome 15 and encompass 4 genes (*NIPA1, NIPA2, CYFIP1* and *TUBGCP5*) within a 500kb region (Figure 1). The BP1-BP2 region is immediately adjacent to the imprinted regions of chromosome 15 that cause Prader-Willi and Angelman syndromes, but the four genes within the BP1-BP2 region are not themselves imprinted. Individuals with this microdeletion have increased risk of a range of neuropsychiatric phenotypes including developmental and speech delays ^11-13^, autism spectrum disorders ^14^, attention deficit disorders (ADD) ^15^ and schizophrenia ^16^. As with other CNVs causing complex phenotypes (for example 22q11.2 deletion, 1q21.1 duplication), penetrance and expressivity shows wide variation. Penetrance of any phenotype with 15q11 BP1-BP2 deletion has been estimated at ∼10-12% ^11, 17^ which is relatively low compared to other CNVs commonly associated with genetic disorders. Taken together, previous studies estimate the prevalence of BP1-BP2 deletion as approximately 0.25% in apparently healthy controls ^14^, with individual studies ranging from 0.19% to 0.47% in cohorts of up to 101,655 individuals ^11, 16-20^.

**Figure 1.**
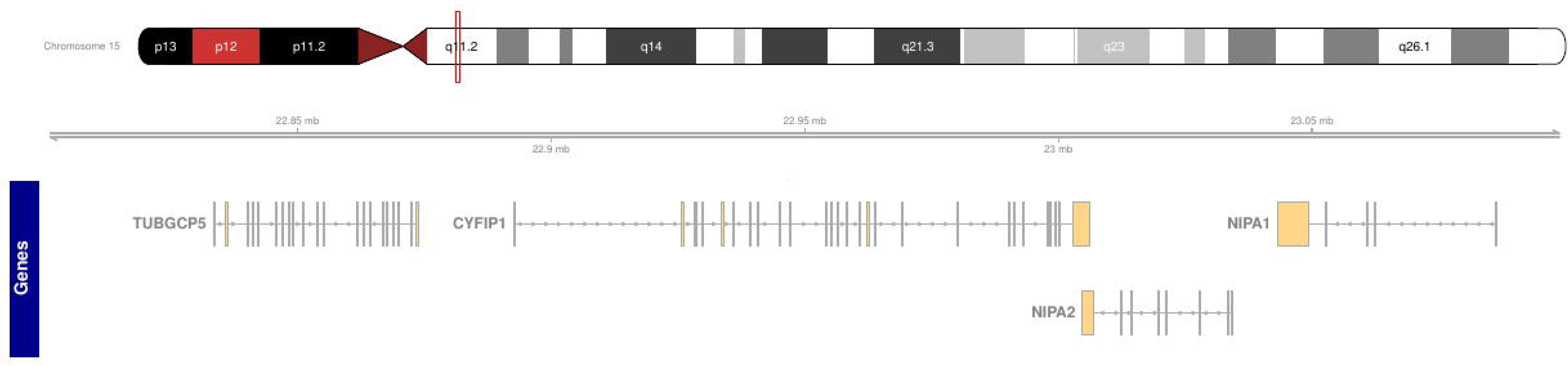
Ideogram of chromosome 15 showing the location of 15q11.2 and the genes between BP1 and BP2.

Genomic deletions are known to be a significant cause of congenital malformations, among which congenital cardiovascular malformations (CVM) have the highest population prevalence ^21-25^. We previously identified twelve heterozygous 15q11.2 BP1-BP2 deletions in a cohort of 2,256 patients with a range of non-syndromic CVM phenotypes, and only 1 in 1,538 controls, inferring association of the deletion with CVM risk (OR = 8.2; p = 0.02) ^26^. Other studies have found the deletion over-represented in patients presenting with intellectual disability who also have CVM ^11, 15, 27, 28^ compared to controls, but data is sparse regarding the association of CVM with this deletion in large populations unselected for disease.

The UK Biobank (UKB), comprising ∼500,000 individuals with genotype data, hospital admission records, baseline cognitive function testing, and a variety of other collected information, provides a large-scale resource for investigating complex genotype-phenotype associations. Here we determine the prevalence of BP1-BP2 deletions within the UK Biobank cohort and explore its association with neurodevelopmental diagnoses, cognitive function and CVM.

## Methods

### CNV calling

B-allele frequency (BAF) and log2ratio (L2R) transformed intensity value files, generated and made available by UKB using the Applied Biosystems UK BiLEVE Axiom and UK Biobank Axiom arrays (n=488,366), were used to generate CNV calls using the PennCNV software ^29^. Samples were grouped for CNV calling in batches of ethnically similar samples based on the ethnicity categories defined by UKB (White, Asian or Asian British, Black or Black British, Chinese, Mixed, Other/Unknown). Ethnically ‘white’ samples were further batched for CNV calling into batches of 10,000 samples due to the large number of individuals in this category. PennCNV was run using GCmodel adjustment ^30^ and, following genome-wide CNV calling, samples with waviness factor (WF) less than -0.03 or greater than 0.03 were excluded (n=301). Samples containing an excess of CNVs (>40) (n=1401) were also removed at this stage. This threshold intentionally retained samples with a relatively high number of CNVs as global CNV burden has previously been found to contribute to the risk of sporadic CVM ^26^.

15q11.2 deletions were identified where a heterozygous deletion spanned at least 95% of the 4 genes between involved (*NIPA1, NIPA2, CYFIP1* and *TUBGCP5*) between the outer-most array probes covering this region (chr15:22,819,338-23,093,090; GRCh37).

### Neuropsychiatric classification

The hospital episode statistics (HES) available in the UKB contains information on hospital admissions for the cohort. It includes primary and secondary diagnoses in the form of ICD9 and ICD10 codes, as well as details of operations and procedures through OPCS-4 codes. Classification of neuropsychiatric conditions was determined using ICD10 codes relating to a subset of ‘mental and behavioural disorders’ (Table S1A) including schizophrenia, bipolar disorders, autism, mental retardation and various neurodevelopmental disorders to classify case individuals. Individuals with any other code in the mental and behavioural classification (Table S1B) were excluded from the control group. This resulted in 3504 case and 483,330 control individuals for neuropsychiatric disorders with CNV calls passing QC thresholds that were then assessed for the presence of BP1-BP2 deletion.

### Cognitive function

UKB participants completed a series of cognitive function tests. We correlated performance in these tests with the presence and absence of the BP1-BP2 deletion. Cognitive function in carriers of a range of neurodevelopmental CNVs has been previously assessed in the first release of UKB (∼150,000 individuals) ^31^. Here, as BP1-BP2 deletions have been shown to associate with developmental delay, autism and behavioural issues previously, and our focus was to investigate the impact of the deletion in people without diagnosed neuropsychiatric conditions, we removed samples from the whole cohort with ICD10 codes present in the HES with specific diagnoses relating to mental and behavioural disorders previously associated with 15q11.2 deletion (Table S1A). In total there were 3,691 samples that matched at least one of these ICD10 codes and were subsequently removed from the cognitive function analysis.

We used results from cognitive function tests where the number of participants was at least 10% of the cohort and the test returned numerical data. Tests for ‘reaction time’, ‘numeric memory’, ‘fluid intelligence’ and ‘pairs matching’ met these criteria with varying amounts of missing data (Table S2). The test for reaction time is based on the card game ‘snap’. A button is pushed by the participant as rapidly as possible when two cards shown on a screen are matching. We used the reported mean time to correctly identify matches over 12 rounds. In the numeric memory test a participant is shown increasingly long numbers (in terms of digits) and asked to recall them after a short period. Here we took the maximum number of digits correctly remembered. In the test for fluid intelligence a participant has 2 minutes to complete as many questions as possible that require logic and reasoning ability, independent of acquired knowledge. The score is the number of correctly answered questions. Finally, the pairs matching test asks participants to memorise the position of matching pairs of cards that are then turned over. We have used the number of incorrect matches in a round as an indication of success in this test.

### CVM classification

Using the classification schema shown in Figure 2, ICD9, ICD10, OPCS-4 and self-reported illness and operation codes were used to classify 2792 UK Biobank samples as having CVM and 472,378 samples as controls in a similar protocol to Saha et al. ^32^. To ensure that CVM-classified individuals were non-syndromic we excluded a number of syndromes with possible links to cardiovascular defects from the cohort (codes listed in Table S3A). Following this, the initial inclusion criterion for case status was HES evidence of ICD9 or ICD10 codes for ‘congenital malformations of the circulatory system’ (Table S3B). Within these, the ICD10 code Q211 can be used for both ‘atrial septal defect’ (ASD) and ‘patent foramen ovale’ (PFO). Since PFO is a normal variant found in up to 25% of the population, we carried out further classification to identify and remove PFOs from the case group. Any sample with a PFO-specific operation code (K165) in addition to a Q211 code was classified as PFO. As PFO is often diagnosed during a stroke workup ^33^ we also used any diagnosis of stroke without atrial fibrillation prior to Q211 diagnosis as an indicator of PFO rather than ASD. This is because ASD is a common cause of atrial fibrillation, which is a risk factor for stroke, whereas PFO (prior to device closure) is not associated with a significantly increased predisposition to atrial fibrillation (all codes in Table S3C). OPCS-4 codes for operations commonly associated with congenital heart defects were also used to classify CVM (Table S3D).

**Figure 2.**
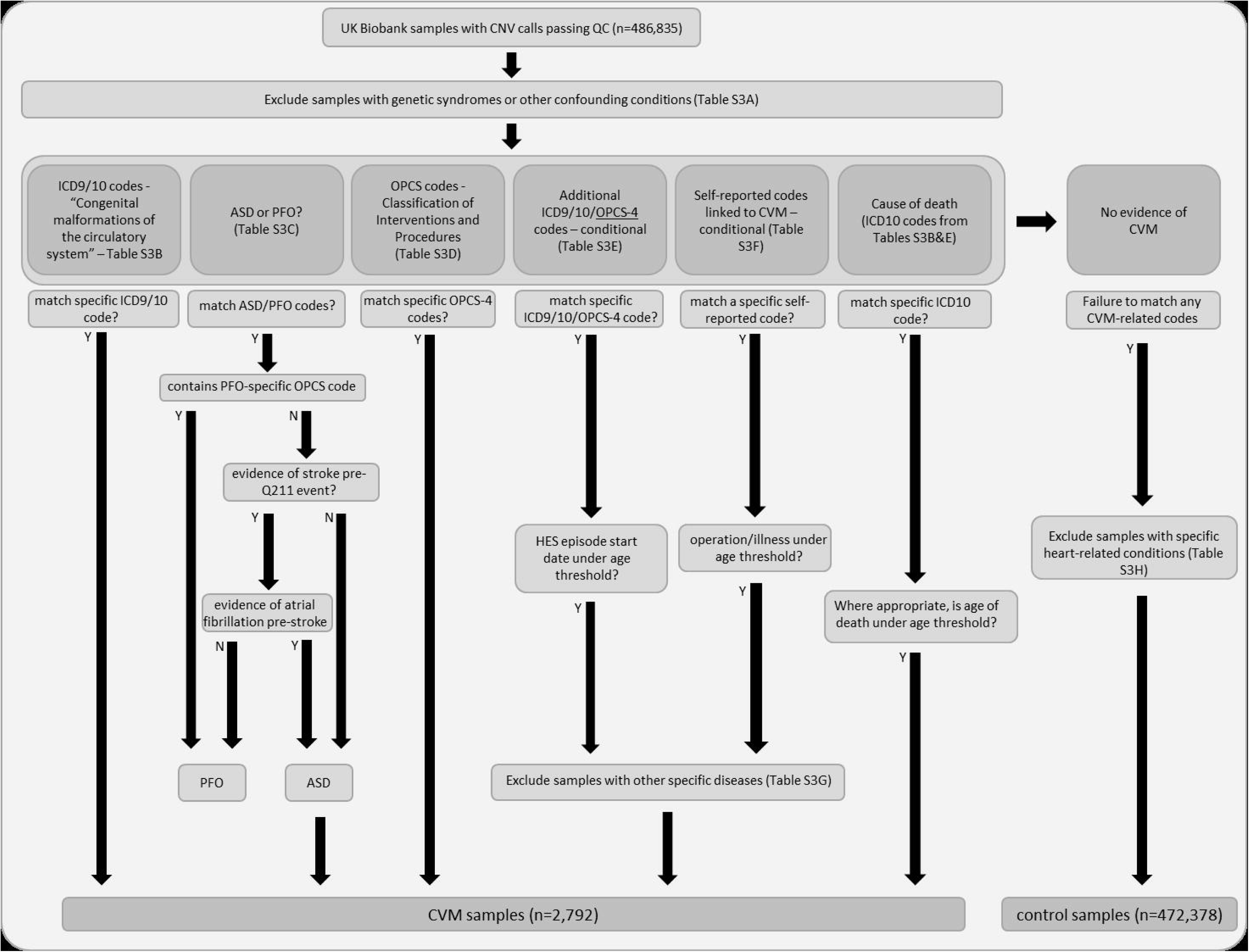
The CVM and control sample classification process on UK Biobank samples using a combination of codes relating to HES data and self-reported information with appropriate thresholds and filtering. Specific codes for each classification step can be found in Table S3.

In addition to codes covering ‘congenital malformation of the circulatory system’, we used additional codes to identify, amongst others, certain defects of congenital origin presenting in later life (Table S3E). Chief among these were codes indicating congenital bicuspid aortic valve (BAV), a condition affecting ∼1% of males and ∼0.33% of females. We included codes for aortic stenosis and aortic insufficiency as well as associated operational codes. We also used self-reported operation and illness codes (full list Table S3F) indicating ‘aortic valve replacement’, ‘aortic stenosis’ and ‘aortic regurgitation’ as evidence of BAV. An age threshold of diagnosis at younger than 65 years was applied to distinguish individuals with aortic valve disease due to BAV from those with age-related degeneration of a trileaflet aortic valve ^34^, as previously implemented by Helle et al. ^35^. Additional filters were applied to exclude any participants with a diagnosis of other conditions (such as bacterial endocarditis) that might cause non-congenital valve defects. The codes and descriptions of these exclusion diagnoses are listed in Table S3G. Individuals with no evidence of CVM phenotypes were classified as controls. Other diagnosed conditions that could potentially have a confounding influence on the classification were excluded from the control set (e.g. PFO-categorised samples, acquired heart defects, samples with evidence of aortic valve disease above the specified age threshold). The full list of excluded codes can be found in Table S3H. Finally, deceased UKB participants with or without CVM classifying ICD codes, obtained by UKB from the UK Office for National Statistics, were classified as CVM cases or controls accordingly.

### Statistical analysis

The frequency of BP1-BP2 deletion in cases of CVM and controls as well as neuropsychiatric disorder cases and controls was compared using chi-squared tests. Associations between the deletion and quantitative measures of cognitive function were assessed using different statistical models dependent upon the distribution of the data, and adjusting for age and sex. Linear regression was used for reaction time (removing values greater than 5x standard deviation from the mean). Poisson regression was fitted for fluid intelligence score. For the pairs matching test where values are heavily skewed towards zero, values were converted to a binary variable (0(0), 1(>0)) and logistic regression fitted on the binary outcome. Assessment of the numeric memory test was again assessed with logistic regression by converting the values into a binary variable (0(<=6), 1(>6)). Association between the deletion and the highest qualification achieved was assessed by combining ‘O levels/GCSEs or equivalent’, ‘CSEs or equivalent’, ‘NVQ or HND or HNC or equivalent’, ‘Other professional qualifications’, ‘None of the above’ and ‘Prefer not to answer’ into a single category and treating this as a baseline group. Further details on UK specific qualifications are available at https://www.gov.uk/what-different-qualification-levels-mean/list-of-qualification-levels. The highest qualification achieved per individual was assessed in terms of ‘College/University degrees’ or ‘A levels/AS levels or equivalent’ and each of these categories compared to the baseline group by fitting a multinomial log-linear model, accounting for age and sex. Fecundity was assessed in males and females separately using Poisson regression fitted to the number of children fathered and number of live births per person respectively. Outliers were removed if greater than 5x standard deviation from the mean. This excluded samples with >8 children fathered in males and >7 live births in females. An interaction test was performed to confirm the differences between male and female subgroups.

## Results

### The prevalence of BP1-BP2 deletion in neuropsychiatric disorder cases

We first assessed the prevalence of BP1-BP2 deletion in individuals diagnosed with a neuropsychiatric disorder. 3504 individuals were found to have a relevant diagnosis in the HES data (see Methods). Of these, 24 carried the deletion, a prevalence of 0.68%, which was different from the control group (OR=1.84 [95% CI 1.23 – 2.75]; p=0.0043) (Table 1). The specific diagnoses of these 24 individuals can be found in Table S4A.

**Table 1.** The number and prevalence of BP1-BP2 deletions in neuropsychiatric cases compared to controls in UK Biobank.

|  | BP1-BP2 del | No BP1-BP2 del | Prevalence |
| --- | --- | --- | --- |
| Non-neuropsychiatric controls | 1808 | 481522 | 0.37% |
| Neuropsychiatric cases | 24 | 3480 | 0.68% |

### BP1-BP2 deletions and cognitive function

After removing participants with relevant neuropsychiatric disorders (See Methods), 483,330 samples remained. Of these, 1808 carried the BP1-BP2 deletion. Figure 3 shows the differences between the two groups in terms of the scores in 4 different cognitive function tests. Table 2 summarises the results of these tests and indicates differences in performance between individuals with and without BP1-BP2 deletion, after adjustment for participant age and sex.

**Table 2.** Summary of the cognitive function tests in the BP1-BP2 deletion group and non-carriers.

| Test | BP1-BP2 del |  |  |  | No BP1-BP2 del |  |  |  | p value |
| --- | --- | --- | --- | --- | --- | --- | --- | --- | --- |
|  | N | Mean | Median | SE | N | Mean | Median | SE |  |
| Reaction time (milliseconds) | 1781 | 574.60 | 551.0 | 2.789 | 475318 | 556.80 | 536.0 | 0.160 | 1.38E-13 |
| Fluid intelligence score | 556 | 5.40 | 5.0 | 0.080 | 158373 | 5.99 | 6.0 | 0.005 | 8.02E-09 |
| Numeric memory | 188 | 6.07 | 6.0 | 0.140 | 50158 | 6.48 | 7.0 | 0.008 | 9.00E-06 |
| Pairs matching | 1808 | 0.61 | 0.0 | 0.030 | 480590 | 0.54 | 0.0 | 0.002 | 2.59E-03 |

**Figure 3.**
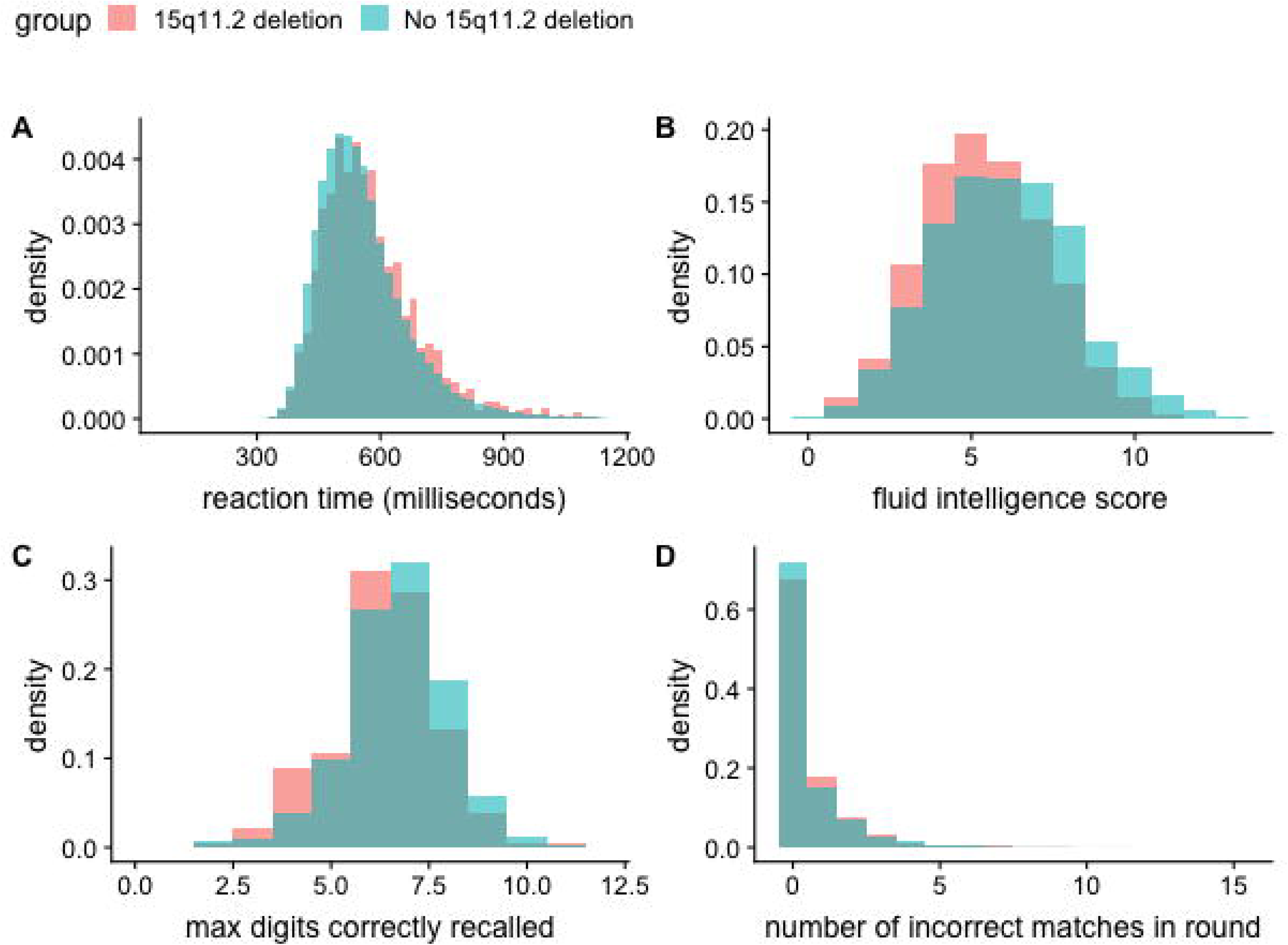
Cognitive function test differences between individuals carrying a 15q11.2 deletion and those without. **A.** Reaction time test. **B.** Fluid intelligence score. **C**. Numeric memory test. **D.** Pairs matching test.

### BP1-BP2 deletions and academic achievement

Carriers of the BP1-BP2 deletion were compared to non-carriers in terms of the highest academic qualifications that they had achieved. Table 3 shows the number of each cohort with ‘College/University degree’ and ‘A levels/AS levels’ as their highest qualification compared to a combined ‘Other categories’ group. We observed lower proportions of deletion carriers obtaining college/university degrees and Advanced and AS level qualifications (typically taken at age 17-18 and required for university or college entrance). 32.8 % of participants without BP1-BP2 deletion had attained a College/University degree, whereas 22.8% of deletion carriers had done so (OR=0.57 [95% CI 0.51-0.64]; p=5.60E-22). Similarly, a lower proportion of participants with BP1-BP2 deletion had attained A/AS levels as their highest educational qualification than participants without the deletion (OR=0.76 [95% CI 0.65-0.88]; p=4.12E-04).

**Table 3.** Academic qualifications achieved by carriers of BP1-BP2 deletion in comparison with non-carriers. ‘College/University degree’ and ‘A levels/AS levels’ highest qualification attainment groups are each compared with a combined ‘Other categories’ group. Estimated odds ratios reflect the chance of the BP1-BP2 deletion group achieving the qualification compared to non-carriers.

| **Highest qualification achieved** | **BP1-BP2 deletion**<br>**frequency** | **No BP1-BP2 deletion**<br>**frequency** | **odds** | **p value** |
| --- | --- | --- | --- | --- |
| College/University degree | 414 | 157988 | 0.57 | 5.60E-22 |
| A levels/AS levels | 185 | 54077 | 0.76 | 4.12E-04 |
| Other categories | 1209 | 268962 | - | - |

### The prevalence of BP1-BP2 deletion in non-syndromic cardiovascular malformation

In total, 2792 participants with genetic data available were identified as having non-syndromic CVM and 472,378 were classified as controls (See Methods). Table 4 shows a list of the most common CVM diagnoses and associated sample numbers in UKB based upon compiling relevant ICD9, ICD10, OPCS-4 and self-reported codes. A full list of these codes along with diagnoses and associated number of individuals can be found in Table S5. Of the participants with CVM, the most common diagnoses relate to the presence of bicuspid aortic valve (BAV). We inferred bicuspid aortic valve from any aortic valve diagnosis (aortic stenosis, regurgitation, unspecified problem or replacement) after childhood but before age 65 in the absence of a diagnosis of infective endocarditis (see Methods).

**Table 4.** The most common CVM diagnoses in UK Biobank.

| **CVM diagnosis** | **Cases** |
| --- | --- |
| Aortic valve replacement (<65 years) | 896 |
| Aortic stenosis (<65 years) | 841 |
| Aortic insufficiency (<65 years) | 594 |
| Atrial septal defect | 409 |
| Malformation of vascular system | 369 |
| Heart surgery (<18 years) | 212 |
| Congenital heart disease - unspecified | 110 |
| Ventricular septal defect | 95 |
| Cardiac septum defect - unspecified | 90 |
| Pulmonary insufficiency | 50 |
| Pulmonary stenosis | 45 |
| Aortic valve issue - unspecified (<65 years) | 36 |
| Coarctation of aorta | 30 |
| Pulmonary artery issue - unspecified | 29 |
| Patent ductus arteriosus | 28 |
| Aortic defect - unspecified | 23 |
| Dextrocardia | 19 |
| Heart block | 17 |
| Tetralogy of Fallot | 17 |
| Aortic atresia | 13 |

In total, 1832 BP1-BP2 deletions were identified in UKB participants with genotyping data available. The CVM and control groups accounted for a total of 1787 of the samples with BP1-BP2 deletion, with 45 deletions present in participants who were excluded according to the criteria above. Of the 2792 non-syndromic CVM samples, 18 carried the BP1-BP2 deletion resulting in an estimated prevalence of 0.64% (Table 5). The specific phenotypes and classifying codes of these cases are summarised in Table S1B. Of 472,378 control samples, 1769 carried the BP1-BP2 deletion resulting in a prevalence of 0.38% (OR=1.73 [95% CI 1.08 – 2.75]; p=0.02995). In keeping with previous observations, no participant was homozygous for the BP1-BP2 deletion.

**Table 5.** The number and prevalence of BP1-BP2 deletions in non-syndromic CVM cases compared to controls in UK Biobank.

|  | BP1-BP2 del | No BP1-BP2 del | Prevalence |
| --- | --- | --- | --- |
| Non-CVM controls | 1769 | 470609 | 0.38% |
| All non-syndromic CVM samples | 18 | 2774 | 0.64% |

Two of the 18 CVM-case samples carrying the BP1-BP2 deletion were also found to have neuropsychiatric-related diagnoses. Those individuals had cardiovascular diagnoses of atrial septal defect and aortic valve disease/replacement; and neuropsychiatric diagnoses of bipolar disorder and scholastic developmental disorders respectively.

### BP1-BP2 deletions and fecundity

Fecundity was assessed in those individuals with and without the BP1-BP2 deletion. Self-reported number of children fathered (for males) and number of live births (for females) in each cohort is shown in Figure 4. A difference was identified in males between those individuals carrying the BP1-BP2 deletion and those without (mean number of children fathered = 1.66 and 1.8 respectively; p=0.00175). This difference was not evident in females (mean number of live births = 1.88 and 1.82 in BP1-BP2 deletion carriers and non-carriers respectively; p=0.18). A test for interaction confirmed the differences between the male and female subgroups (p-interaction=5.46E-04).

**Figure 4.**
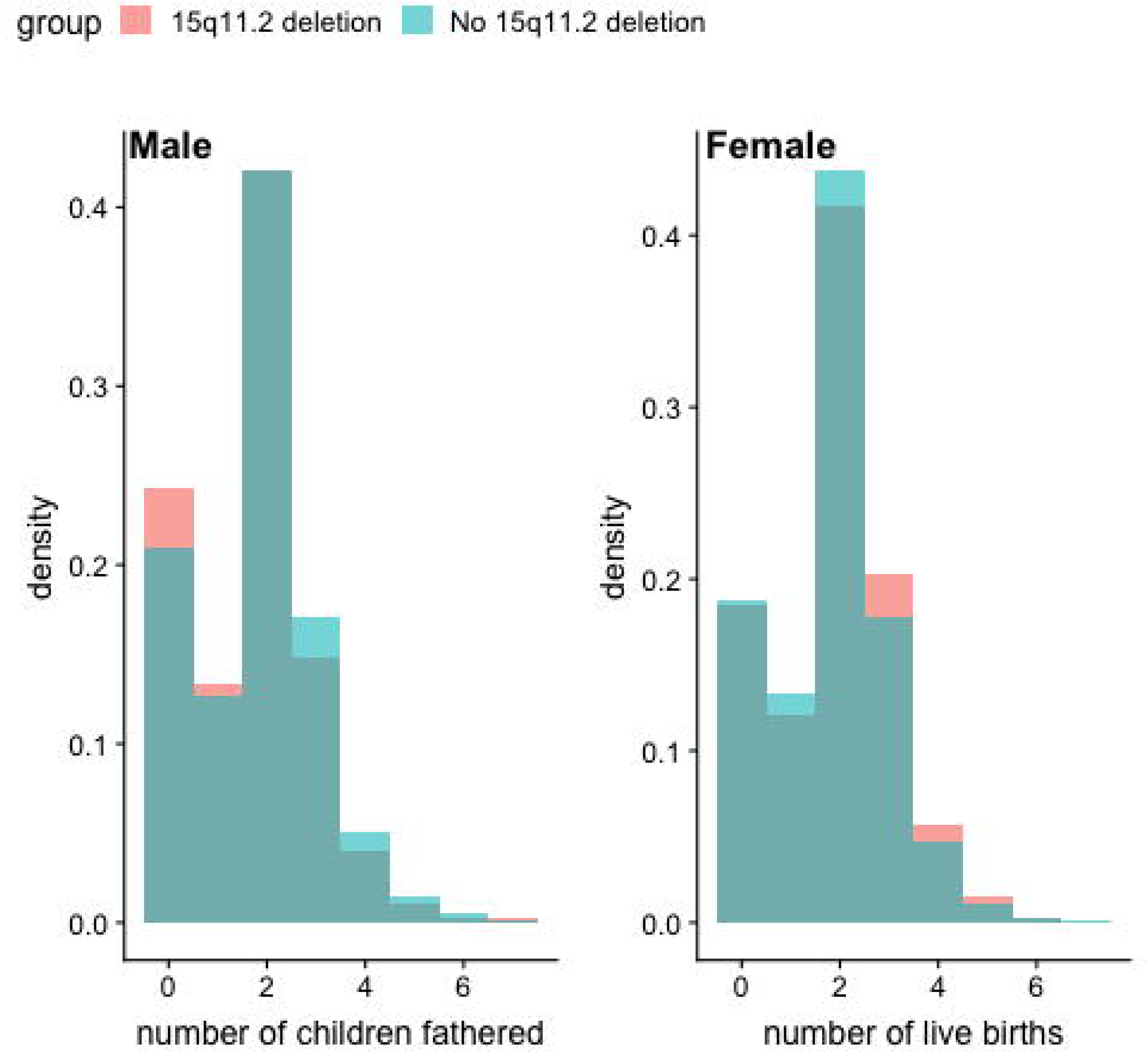
Fecundity measured as self-reported number of children fathered (by males) and number of live births (by females) in individuals with and without the BP1-BP2 deletion.

## Discussion

The 15q11.2 (BP1-BP2) deletion (sometimes referred to as the Burnside-Butler syndrome susceptibility locus) has previously been associated with phenotypes including developmental delay, autism, schizophrenia and CVM; the great majority of the evidence regarding the deletion thus far originates from cohorts specifically selected for one or other of those phenotypes. We estimated the prevalence of the deletion and its associations in the UK Biobank cohort. We confirmed the association of the deletion with neuropsychiatric disorders (OR=1.84 [95% CI 1.23 – 2.75]; p=0.0043); and with CVM (OR=1.73 [95% CI 1.08 – 2.75]; p=0.03).

Estimates of the prevalence of BP1-BP2 deletion have varied widely in the literature. The largest previous study, by Stefansson et al. ^20^ showed a general population prevalence of 0.23% (95% CI 0.21-0.26), in 101,655 Icelandic subjects of whom 241 carried the BP1-BP2 deletion. Given differences in recruitment strategies, genotyping methodologies and populations, direct comparison between this figure and our estimate of 0.37% (95% CI 0.36-0.39) is not straightforward; however, they are broadly consistent with each other. The previous study by Stefansson had also shown nominally reduced fecundity among 172 carriers of the BP1-BP2 deletion who were over 45 years of age. In UKB we observed reduced fecundity in males carrying this deletion (p=0.00175) (Figure 4), supporting these findings in a much larger cohort. Stefansson et al. concluded that the neuropsychiatric manifestations of the BP1-BP2 deletion in carriers without schizophrenia were most evident on dyslexia and dyscalculia ^20, 36^. The present study had a less extensive range of cognitive function tests, but with larger numbers of deletion carriers we observed a more general effect on cognitive function among carriers who had not been diagnosed with a neuropsychiatric disorder. Additionally, we showed sizeable associations between the deletion and lower educational attainment. Deletion carriers were considerably less likely to obtain Advanced level qualifications, which in the UK system are required for University entrance; and whereas about one-third of UKB participants without BP1-BP2 deletion had a University or College degree, less than one-quarter of deletion carriers had this as their highest educational qualification.

We confirmed the association of BP1-BP2 deletion with CVM, which has been observed in some but not all previous studies. The first study linking BP1-BP2 deletions with non-syndromic CVM showed an odds ratio of 8.2 (P=0.02); the risk associated with the deletion was much smaller in this study. Of note, there are major differences in the range of phenotypes in both studies, with a much milder range of CVM phenotypes predominating in UKB. Indeed, CVM phenotypes occurred at lower rates than anticipated in UKB, which is known to exhibit a “healthy cohort effect” in multiple domains. Most UKB participants with CVMs had bicuspid aortic valve, which was excluded from the hypothesis-generating study; moreover, the prevalence of BAV diagnosed from the HES data in UKB was significantly lower than anticipated, indicating misclassification bias could affect the risk estimate. Larger cohorts of homogeneous phenotypes will be required in order to establish the role of the deletion in specific CVM phenotypes. In future studies, availability of electronic health records from the General Practitioners of UKB participants, may mitigate any misclassification bias in respect of CVMs.

The clinical and genetic aspects of BP1-BP2 deletion have been recently reviewed ^37^. Four genes, *TUBGCP5, CYFIP1, NIPA1*, and *NIPA2*, are located in the deleted region. The deletion is recognised as a susceptibility locus for presentations with intellectual impairment, autism, schizophrenia, and epilepsy with a low penetrance estimated at around 10%. The effects of hemizygosity at each individual gene in the region have not been systematically described in mouse models, nor has any mouse model of the entire deletion been reported as yet. Therefore, the mechanisms whereby the deletion causes its associated phenotypes, and the interaction between hemizygosity for the region and other genetic and environmental factors to result in the wide variety of phenotypes described, remain to be elucidated.

In conclusion, we have explored the phenotypic associations of the BP1-BP2 deletion in the largest cohort studied to date. We demonstrate an association with the less severe CVM phenotypes prevalent in the UKB cohort, albeit of small magnitude, as well as confirming the association with neuropsychiatric disorders. We have found broad general effects on cognitive function in carriers of the deletion and evidence of disparity in academic qualifications that have been achieved by carriers compared to non-carriers. Given reduced cognitive function in non-diagnosed carriers, these findings highlight important clinical and societal implications of the BP1-BP2 deletion, and are of potential use in deciding appropriate criteria for clinical genetic testing.

## Conflict of Interest

none declared

## Supplementary material

**Table S1. A**. List of ICD10 codes used to classify individuals as having had phenotypically relevant neuropsychiatric disorders. **B**. Additional ICD10 codes for ‘mental or behavioural disorders’ that were used to exclude individuals from the control group for neuropsychiatric disorders.

**Table S2**. Amount of ‘missing’ data in the cognitive function tests.

**Table S3.** Lists of ICD9, ICD10 and OPCS4 codes used as inclusion and exclusion criteria for classifying the UK Biobank cohort as non-syndromic CVM samples or controls.

**Table S4**. Phenotypes of UK Biobank participants classified as having **A**. neuropsychiatric diagnoses and a BP1-BP2 deletion. **B**. CVM diagnoses and a BP-BP2 deletion.

**Table S5**. A list of CVM-classifying codes and the number of samples in UK Biobank with these classifications.

